# An RNA guanine quadruplex regulated pathway to TRAIL-sensitization by DDX21

**DOI:** 10.1101/588798

**Authors:** Ewan K.S. McRae, Steven J. Dupas, Evan P. Booy, Ramanaguru S. Piragasam, Richard P. Fahlman, Sean A. McKenna

## Abstract

DDX21 is a newly discovered RNA G-quadruplex (rG4) binding protein with no known biological rG4 targets. In this study we identified 26 proteins that are expressed at significantly different levels in cells expressing wild type DDX21 relative to an rG4 binding deficient DDX21 (M4). From this list we validate MAGED2 as a protein that is regulated by DDX21 through rG4 in its 5’UTR. MAGED2 protein levels, but not mRNA levels, are reduced by half in cells expressing only DDX21 M4. MAGED2 has a repressive effect on TRAIL-R2 expression that is relieved under these conditions, resulting in elevated TRAIL-R2 mRNA and protein in cells expressing only DDX21 M4, and rendering previously resistant cells sensitive to TRAIL mediated apoptosis. Our work identifies the role of DDX21 in regulation at the translational level through biologically relevant rG4 and shows that MAGED2 protein levels are regulated, at least in part, by a rG4 forming potential in their 5’UTRs.

## Introduction

DDX21 is an RNA helicase protein with a diverse set of biological functions. It plays important roles in ribosomal RNA biogenesis^1,2^ by coordinating Pol I transcription and association of late acting snoRNAs with pre-ribosomal complexes^3,4^. DDX21 has also been implicated in double stranded RNA sensing and anti-viral response^5–8^ as well as epigenetic silencing of genes^9^. Abnormal levels of DDX21 protein has been observed in colorectal^10^ and breast cancers^11^ (where it has been correlated with disease free survival, possibly mediated by regulating c-Jun activity and rRNA biogenesis^12,13^) as well as in schizophrenic patients, making it a potential therapeutic target. Recently, we have shown that DDX21 can bind and unwind guanine-quadruplexes (rG4s) and affect the translation of a reporter construct with a rG4 in its 5’ UTR^14^. This activity is mediated by an evolutionarily conserved region repeat region (F/PRGQR) in its C-terminus that preferentially interacts with the backbone of RNA rG4s^15^. It is unlikely that such an activity would exist and persist in an enzyme if there were no biological need for it. Recent iCLIP data using anti-DDX21 antibodies showed that ∼35% of DDX21 bound RNA is mRNA^16^, yet no rG4 targets of DDX21 are currently known.

RNA rG4s are four stranded structures of RNA that form in regions rich in guanines. When four guanines align in a plane they can form an extended hydrogen bonding network called a guanine tetrad. Successive guanine tetrads can pi stack on top of one another to form a stable rG4 structure. The canonical rG4 forming sequence is GGG(N_1-7_)GGG(N_1-7_)GGG(N_1-7_)GGG, where (N_1-7_) represents a short loop of 1-7 nucleotides. However, longer loops as well as interrupted G-tracts have also been shown to still allow for rG4 formation. Canonical rG4s^17^ as well as irregular rG4s^18^ present in mRNA have been shown to modulate translation of the mRNA, either through stalling of scanning pre-initiation complexes on 5’ UTRs^17^, by alternative splicing mediated by rG4s^19–21^, or direction of miRNA binding to 3’ UTRs^22,23^. A transcriptome-wide sequencing approach that exploits the ability of RNA rG4s to cause reverse transcriptase stop (rG4-seq) has identified thousands of putative quadruplexes, many of which are non-canonical rG4s that would likely be missed by bioinformatic searches^24^.

In our current study we use proteomic mass spectrometry (MS) to compare the levels of proteins in cell populations that are expressing wild type DDX21 or a DDX21 mutant with impaired rG4 binding and unwinding abilities (DDX21 M4). We compare this list of candidate proteins to the rG4-seq database, finding a significant enrichment for rG4 containing mRNA in our list of candidate proteins, and confirm the interaction of DDX21 with the mRNA of candidate proteins by RIP-qPCR experiments. One of the promising candidate proteins, Melanoma associated antigen D2 (MAGED2) protein abundance is significantly reduced in the cells expressing DDX21 M4 and not the WT DDX21. MAGED2 has been implicated in cancer progression and resistance to treatment of human breast cancers, specifically, MAGED2 acts as a transcriptional repressor of tumour necrosis factor-related apoptosis-inducing ligand (TRAIL) death receptor 2 (TRAIL-R2) mRNA^25,26^. For decades TRAIL has been studied as a potential natural anti-tumoral protein with the ability to selectively trigger cell death in cancer cells through interactions with surface expressed receptors DR4 and DR5 (TRAIL-R1 and TRAIL-R2)^27^. TRAIL resistance occurs in breast cancer cells when the death receptors DR4 and DR5 (TRAIL-R2) are no longer expressed on the cell surface^28^. TRAIL sensitization is emerging as a novel therapeutic target for treating resistance breast cancers^29,30^. We confirm by qPCR and Western blot that regulation of MAGED2 by DDX21 also affects the downstream target of MAGED2, TRAIL-R2, and imparts a sensitivity to TRAIL mediated apoptosis.

To further investigate the link between rG4 and MAGED2 we synthesized a mutant version of the MAGED2 5’ untranslated region (UTR) (where the G4 is predicted to be) with G to C mutations disrupting all runs of 3 or more consecutive guanines. A side by side comparison of the effect of wild-type and mutant 5’ UTR in a luciferase assay reveals a loss of regulatory effect with the mutant UTR. We provide direct evidence of rG4 formation in the *in vitro* transcribed wild-type 5’ UTR using an RNA G4 specific fluorescent probe, Thioflavin T (ThT), and a modified reverse transcription stop method that demonstrates rG4 unwinding by DDX21. Together these results indicate that DDX21 regulates translation of MAGED2 mRNA in a rG4 dependent manner and contributes to sensitization to TRAIL mediated apoptosis.

## Results

### Identifying putative G4 targets of DDX21

To look for potential G4 targets of DDX21 we wanted to compare cell populations expressing a wildtype and a mutant (M4) DDX21 with impaired G4 binding. The M4 mutant contains a PRGQR to YEGIQ mutation in the C-terminal repeat region of DDX21 that has been previously shown to disrupt rG4 binding and unwinding activities^14^. Differences between the wildtype DDX21 recovery (WT) and M4 DDX21 recovery experiments should be due to differences in DDX21 G4 interactions. Briefly, HEK293T cells were first transfected vector expressing either WT or M4 DDX21. Following this, endogenous DDX21 was depleted by siRNA knock-down. Using label-free whole-cell proteome analysis we identified 26 candidate proteins (Figure 1A) whose expression level was significantly different between the M4 and WT samples. The data in Figure 1A is represented as the ratio of the amount of a protein detected in the M4 vs WT samples from biological triplicate analysis, this allows us to visualize a fold change in protein level in the absence of DDX21’s G4 binding activity. Proteins with large positive values like XPO6 have were detected in greater abundance in the presence of M4 DDX21, indicating that the G4 interaction may be impeding protein expression. Whereas proteins with large negative values like CNOT3 have significantly decreased abundance in the presence of M4 DDX21, indicating that a G4 interaction with DDX21 could be promoting expression of the protein. The efficiency of endogenous DDX21 knock-down was confirmed by MS where DDX21 was undetected in 2 of 3 knock-down samples and 10-fold less than negative control in the third replicate. The exogenous DDX21 (WT and M4) from the recovery samples was on average 10-fold more abundant than the DDX21 present in negative control samples. Efficiency of knock-down and recovery of DDX21 was also assessed by western blot (Figure 1B).

**Figure 1.**
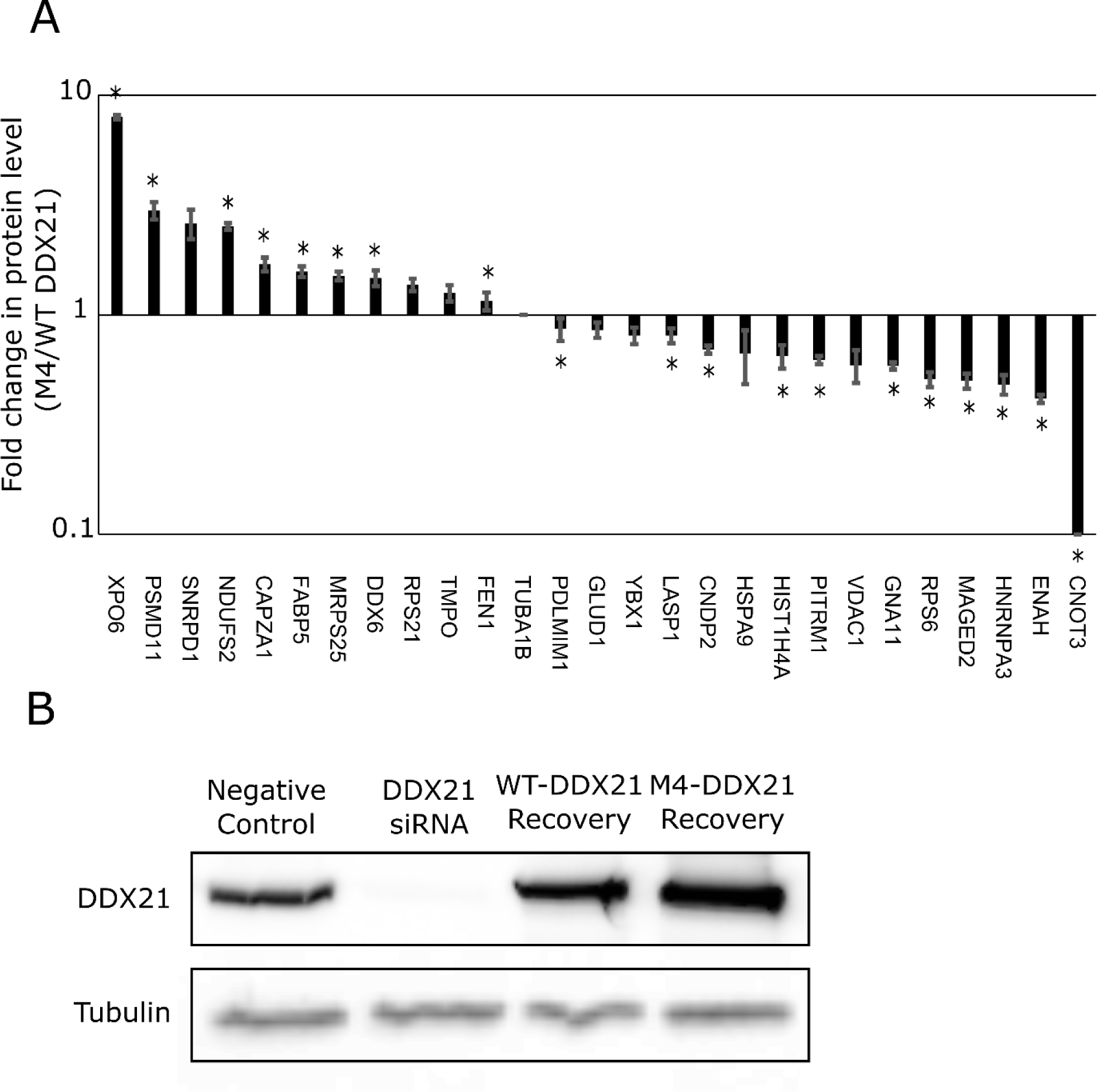
(A) Fold difference in protein levels by MS between HEK293T cells expressing M4 DDX21 over WT DDX21. Two-way Anova tests determined the significance of the difference, error bars represent the standard deviation between 3 biological replicates. (B) Representative Western blot showing DDX21 levels between samples.

To determine if the change in protein levels could be due to altered mRNA level we performed qPCR on RNA extracts from the samples (Figure S1). While some mRNA levels were found to be significantly different than negative control samples (PSMD11, HNRNPA3, TMPO, CAPZA, HIST1H4A & CNOT3), there was no significant difference observed between WT and M4 recovery samples, indicating that the altered mRNA levels are likely artifactual. Since the observed differences between WT and M4 protein levels could not be explained by differences in mRNA abundance, the remaining reasonable explanations are altered translation efficiency or protein stability.

From the list of 26 candidate proteins, 20 (76.9%) were shown to have quadruplex forming potential in their mRNA by rG4-seq experiments^24^. Comparatively, of the 17,622 unique transcripts mapped in the rG4-seq experiment only 37.5% showed evidence of RNA rG4 formation. RIP-qPCR experiments using anti DDX21 antibodies showed significant enrichment (3 standard deviations above GAPDH) for 6 of the candidate proteins’ mRNA (Figure 2). Of these 6 genes, 3 had predicted rG4s in the 5’UTR and 3 had predicted rG4s in the coding sequence (CDS). Interestingly, this distribution is quite the opposite of what was observed in the rG4-seq database in general, they report approximately 62% of the rG4s found to be in the 3’UTR and the remainder to be 16% 5’UTR and 20.6% CDS.

**Figure 2.**
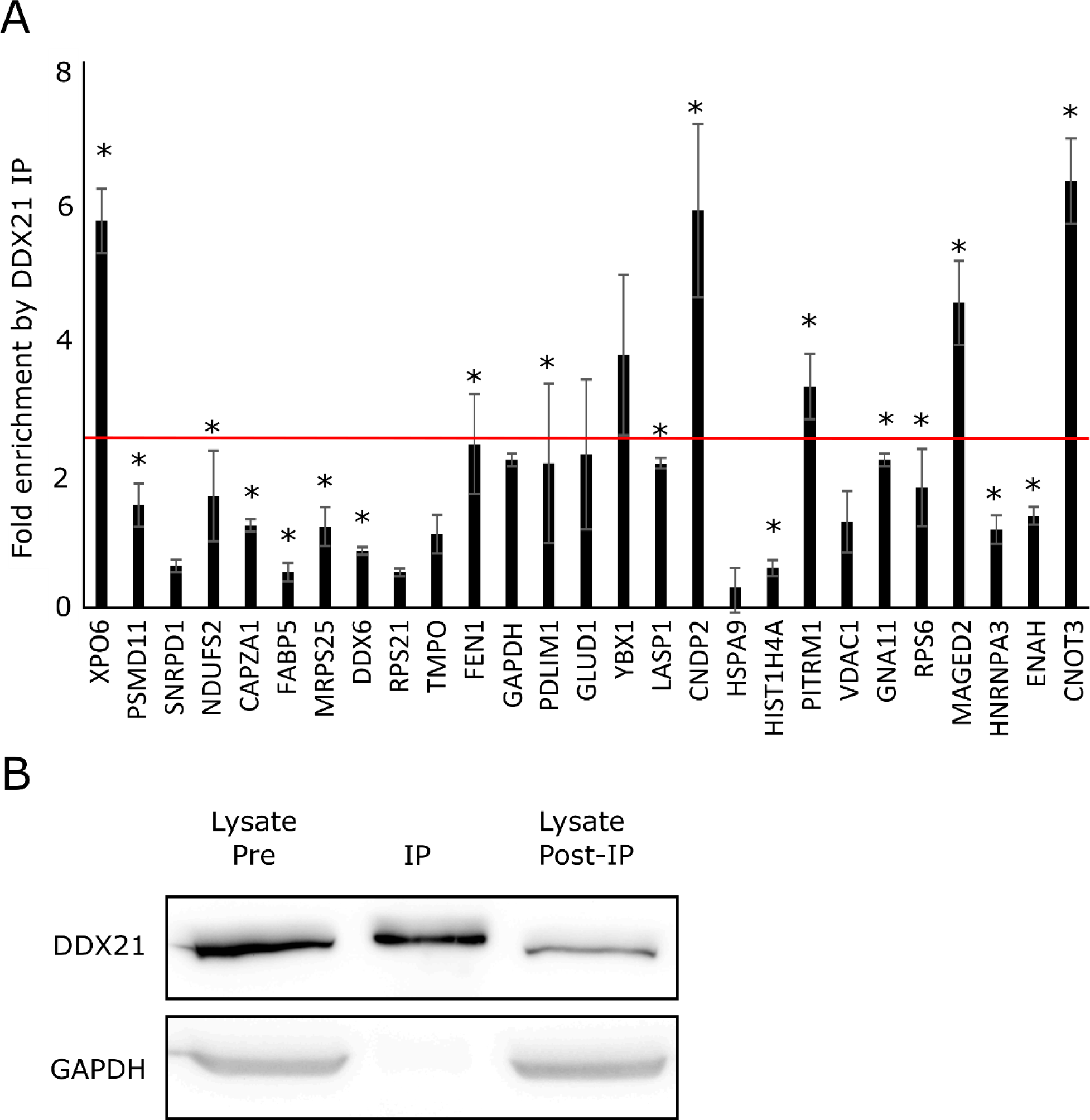
(A) RT-qPCR data for DDX21 RNA IP from HEK293T cell lysate comparing Ct values of DDX21 IP sample to the RNA extracted from the pre-IP lysate. The red line represents a 3SD difference in enrichment above GAPDH, error bars represent the standard deviation between 3 independent IP replicates. (B) Western blot showing DDX21 present in the lysate, IP, and to a lesser extent in the post-IP lysate.

The large enrichment of proteins with rG4 containing mRNA in our list indicates there is good potential for direct regulatory effects to be discovered. The DDX21-IP allows us to further narrow our search by only focusing on the mRNA that are preferentially co-precipitated with DDX21.

### Validating DDX21 targets by Western Blot

We chose to validate the 3 proteins with predicted rG4s in their 5’UTR by Western blot because of the ease of integration into of their G4 containing regions into an existing assay for 5’UTR mediated translational regulation. The two extremes from Figure 1, XPO6 and CNOT3 are such due to XPO6 being undetected in two of three WT DDX21 recovery MS samples and CNOT3 being undetected in all M4 DDX21 recovery MS samples. However, we were able to detect both of these proteins in all samples by Western blot. Western blots (Figure 3) for XPO6 show significant signal in the WT-DDX21 recovery sample and no apparent difference between the other samples. CNOT3 Western blots show increased signal from the WT DDX21 samples compared to all other conditions, but no or marginal difference between the negative control, knock-down or M4 samples. MAGED2 Western blots have decreased signal in the knock-down and M4 samples compared to negative control and WT, consistent with the effect observed by MS. Based on these results we chose to further investigate the role of DDX21 and it’s rG4 binding domain in MAGED2 regulation.

**Figure 3.**
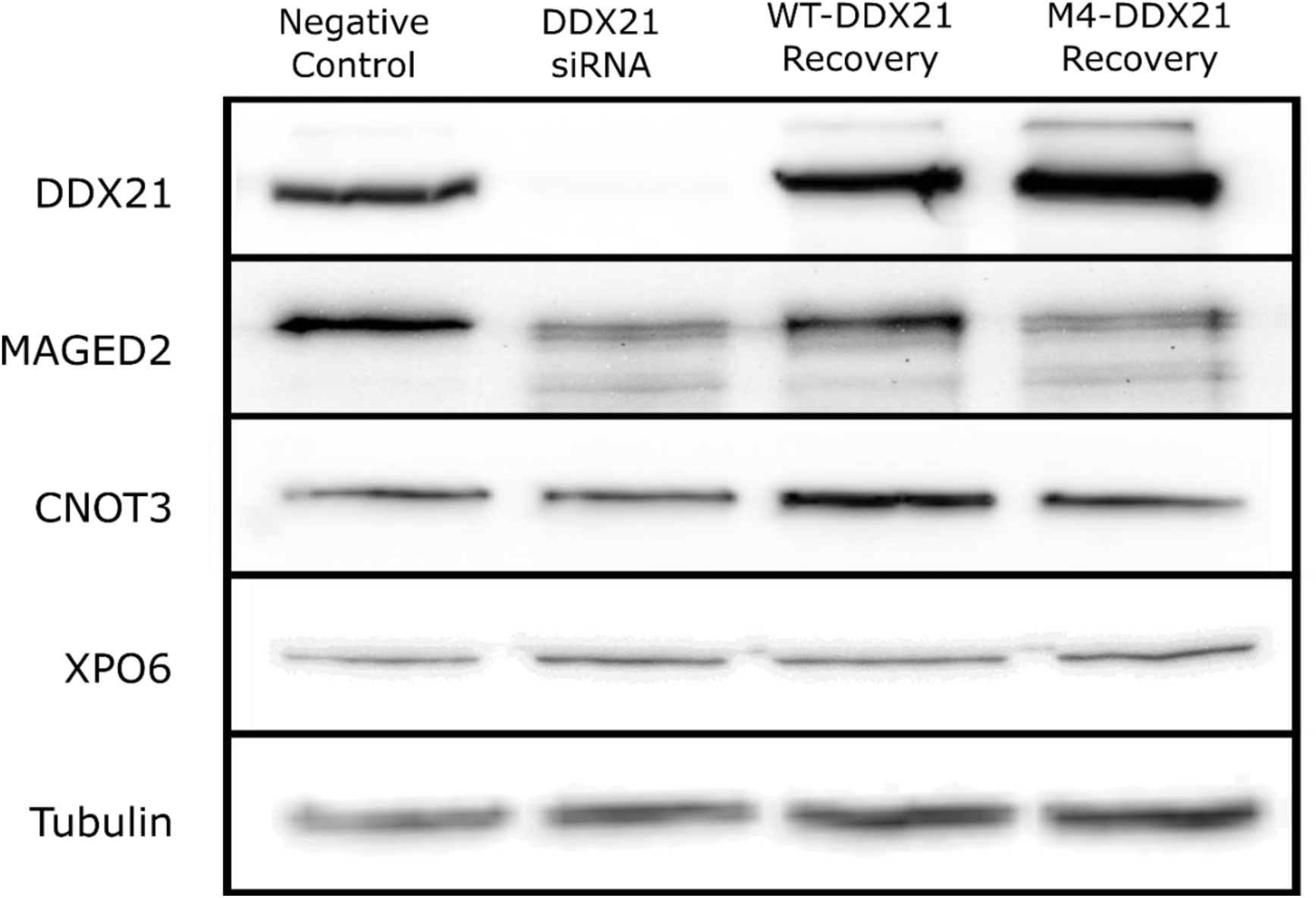
Validation of the changes in protein level observed by mass spectrometry between negative control, DDX21 knock-down and WT or M4-DDX21 recovery samples by Western blot. DDX21 knock-down and recovery (top) affects the signal from the MAGED2 antibody, but not the antibodies for CNOT3, XPO6 or Tubulin.

### DDX21 regulates TRAIL-R2 expression and protects cells from TRAIL mediated apoptosis

MAGED2 has been shown to regulate TRAIL-R2 protein and mRNA levels in a p53 dependent manner^25,26,31^. To test whether the changes in MAGED2 level from DDX21 knock-down or M4 mutant could affect TRAIL-R2 we performed qPCR on RNA extracted from our knock-down recovery experiments. The RNA from the HEK293T cells showed no difference in mRNA levels of p53, MAGED2 or TRAIL-R2 (Figure S1). Since this pathway has been shown to be p53 dependent, and HEK293T have p53 suppressed by the SV40 large T antigen^32^, we repeated the experiment with MCF-7 cells that have wild-type p53^33^. In MCF-7 cells we saw an increase in TRAIL-R2 mRNA levels upon DDX21 knock-down, but not p53 or MAGED2 (Figure 4A). Furthermore, Western blots confirm a change in protein levels of TRAIL-R2 in response to DDX21 activity (Figure 4B). Next, we investigated the TRAIL sensitivity of our four conditions, 72 hours after knock-down of DDX21 MCF-7 cells were treated with TRAIL protein and left overnight. To detect TRAIL mediated apoptosis, we used fluorescent Annexin V to stain cells undergoing apoptosis and counted fluorescent cells by flow cytometry (Figure 4C). The DDX21 knock-down and M4-DDX21 recovery samples were significantly sensitized to TRAIL mediated apoptosis with ∼20% of the counted cells showed staining by Annexin V. Comparatively the negative control treated cells and the WT-DDX21 recovery had only 0.19 ± 0.02% and 0.52 ± 0.18% of cells stained with Annexin V. These results demonstrate that regulation of MAGED2 by DDX21 is sufficient to affect the TRAIL-sensitization of cells by overexpression of TRAIL-R2.

**Figure 4.**
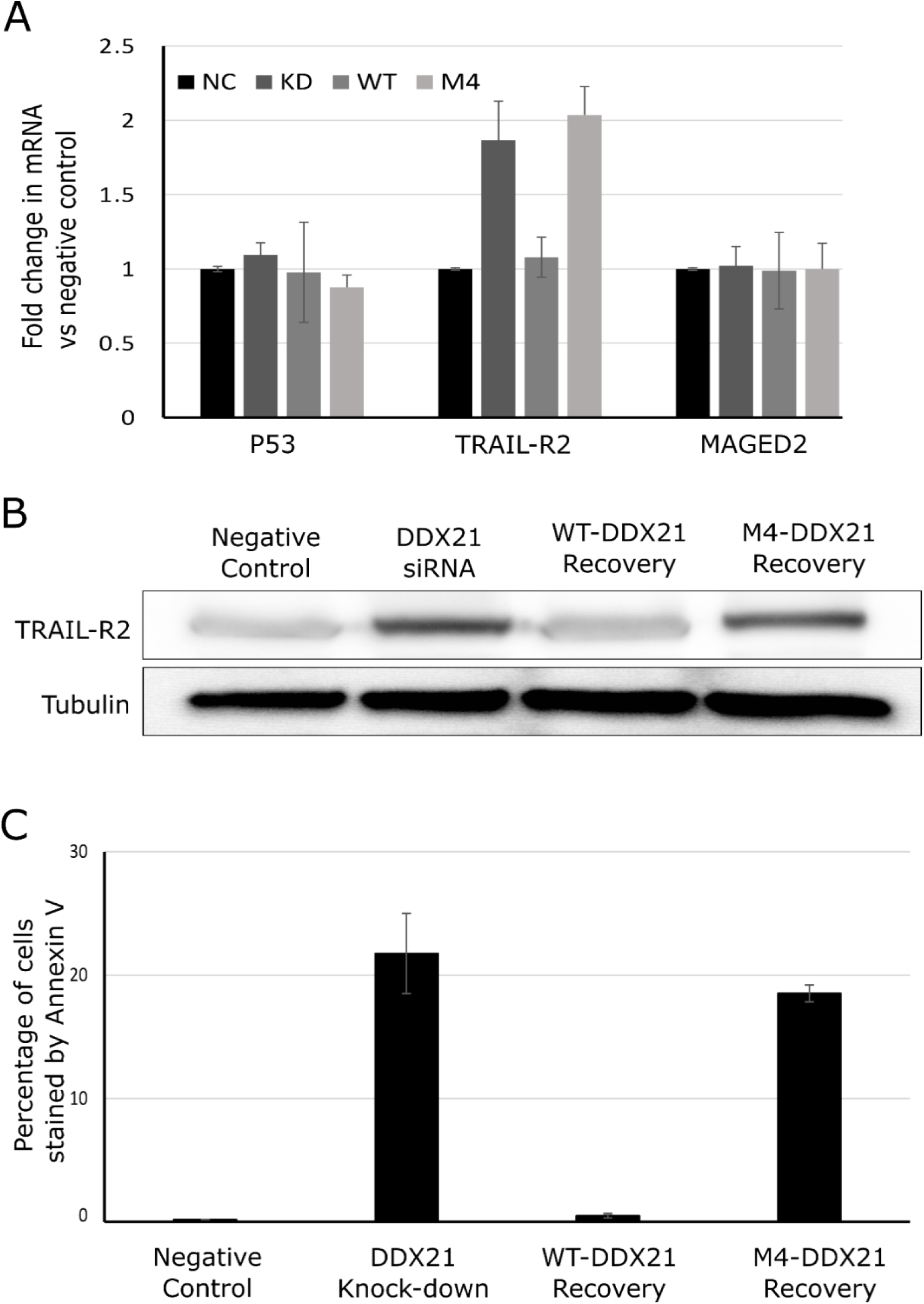
Assessing the effects of DDX21 knock-down, WT and M4 recovery on MCF-7 cells by: (A) RT-qPCR for MAGED2 mRNA and its downstream targets TRAIL-R2 and P53. (B) Western blot for TRAIL-R2 levels. (C) Flow cytometry analysis of Annexin V staining post TRAIL treatment to detect apoptosis. Error bars represent the standard deviation between 3 biological replicates

### MAGED2 TV2 is most enriched transcript variant by RIP-rt-qPCR

The failure of the M4-DDX21 mutant to recover to the wild-type MAGED2 level is indicative of the involvement of rG4s in the regulation. MAGED2 has three alternative splice variants that result in different 5’UTR, but with no change to protein coding sequence. Though the variants share a conserved 31 nucleotides before the start codon, their 5’ regions vary in length and sequence from 125nt for TV1, 233nt for TV2 and 193nt for TV3. All transcript variants have high rG4 forming potential, TV1 has 5 runs of 3 consecutive guanines, TV2 has 9 runs of 3 consecutive guanines and TV3 has 8 runs of 3 consecutive guanines. To decipher which of these transcript variants could be contributing to regulation of MAGED2 we first returned to our DDX21-RNA-IP samples and looked for differences in enrichment between primer sets specific for individual transcript variants (Figure 5A). While a primer set that doesn’t discriminate between MAGED2 transcript variants is enriched 4.6-fold, TV1 and TV3 were only enriched 1.7 and 2.2-fold, respectively. TV2 has the highest fold enrichment with a 5.5-fold change, indicating it is preferentially interacting with DDX21 in cell lysate.

**Figure 5.**
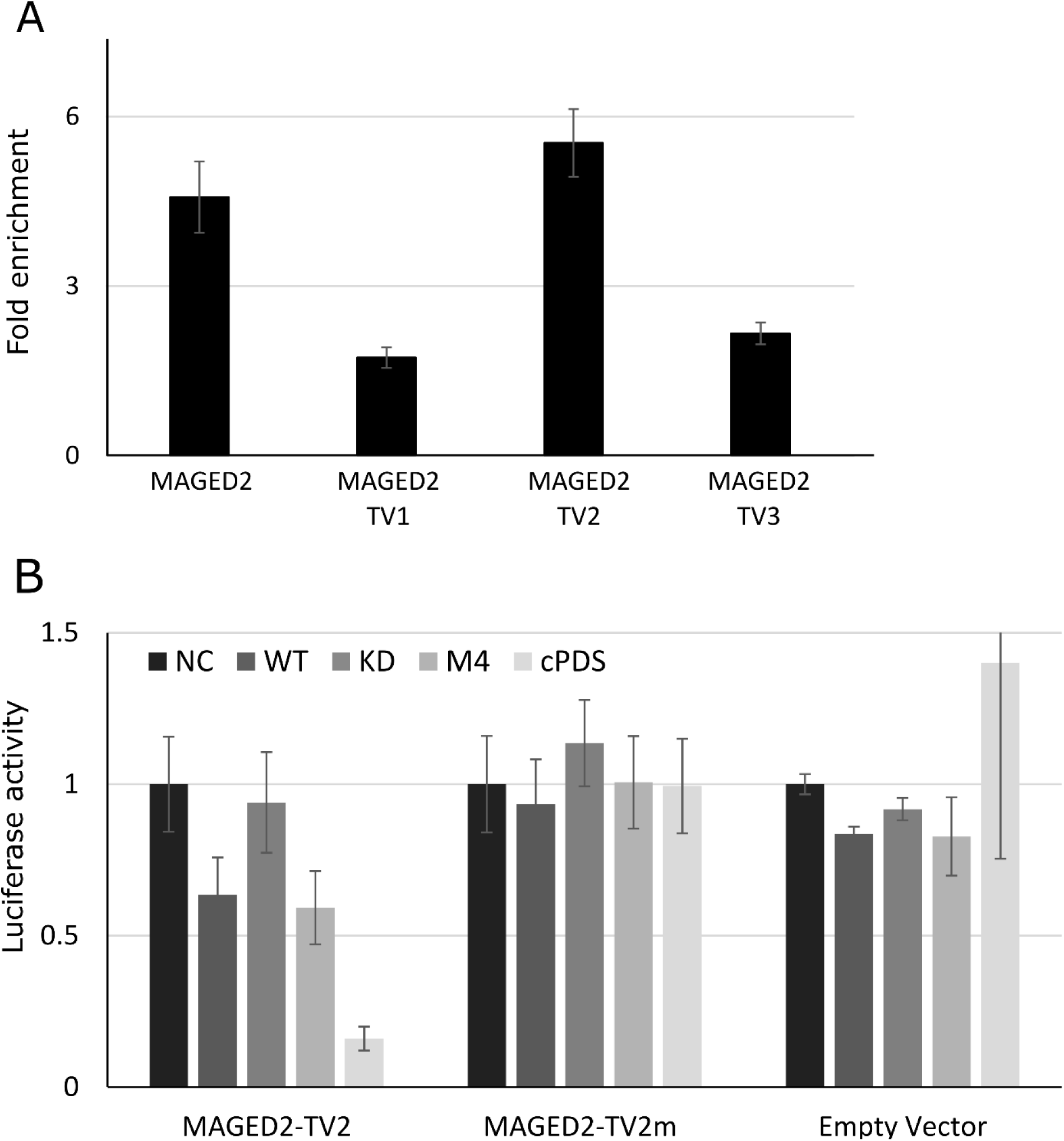
(A) DDX21 RNA IP enriches MAGED2 TV2 preferentially to TV1 and TV3 by RT-qPCR. (B) Looking at the effect of DDX21 status and rG4 stabilizer cPDS on luciferase activity under control of MAGED2 TV2, TV2m and no 5’ UTRs. Error bars represent the standard deviation between 3 biological replicates.

### MAGED2 5’UTR Luciferase Assays

As the strongest candidate for DDX21 interaction we site specifically mutated all 9 runs of guanines from TV2 so that there are never more than two consecutive guanines (TV2m). TV2 and TV2m were cloned into the dual-luciferase vector, PsiCheck-2, immediately 5’ of the hRLuc gene such that they would be transcribed as a 5’UTR. A common criticism of these assays is that the efficient transcription from the promotor on the luciferase vector results in levels of hRLuc mRNA that far exceed the levels of mRNA of the endogenous mRNA. In the context of G4 regulation, the increased number of G4s could exceed the capacity of the cellular G4 helicase proteins to unwind them^34^. In order to mitigate this effect, we truncated the CMV promotor in front of our 5’UTR to reduce the level of reporter mRNA being made. HEK293T cells were treated as they were for the MS experiments and at 48 hours post endogenous DDX21 knock-down they were transfected with the dual-luciferase reporter vector.

In the experiments with the wild type TV2 5’ UTR, DDX21 siRNA knock-down significantly (P<0.05) reduced the luciferase activity from the levels observed in the negative control cells (Figure 5B). This effect was rescued by overexpression of the WT DDX21 but not M4. The luciferase activity from the M4 samples was also significantly reduced compared to negative control and WT samples, but not significantly different than the DDX21 knock-down samples. In the experiments with TV2m or an “empty vector” (no 5’ UTR added), no significant change in luciferase activity was detected between NC, KD, WT or M4 samples.

To begin probing whether the MAGED2 TVs form rG4, we introduced an RNA rG4 stabilizing ligand carboxypyridostatin (cPDS) into the luciferase assays. Cells that were otherwise untreated were treated with cPDS immediately prior to transfection of the reporter vector. Compared to the samples that were not treated with cPDS, the treated samples had only 15% of the luciferase activity from the vector with the intact TV2 5’UTR. Whereas, the TV2m reporter construct showed no change in luciferase expression in the presence of cPDS. The empty vector samples were similarly unaffected by cPDS with the exception of one replicate having increased luciferase activity, adding significant error to the average luciferase activity.

### In Vitro evidence of rG4 formation

To further validate rG4 formation the TV2 and TV2m were *in vitro* transcribed from the PsiCheck-2 vector and purified by size exclusion chromatography. To confirm the possibility of direct interactions with DDX21 and to ascertain any possible specificity of DDX21, electrophoretic mobility shift assays (EMSA) were performed (Figure S2) with recombinant purified DDX21 from *E.coli*. At the highest DDX21 concentration (1000nM) both RNA completely shifted to higher molecular weight complexes. Densitometric analysis using the FluorChem Q software allowed for fitting of the EMSA to a binding model (Table 1). Recombinant purified DDX21 showed marginally higher affinity for TV2 vs TV2m (100nm vs 300nm)

TV2 and TV2m were assayed for rG4 formation by ThT, a fluorescent probe for RNA rG4 formation^35,36^. When ThT is bound to rG4 it exhibits enhanced fluorescence. rG4 conformations are stabilized by monovalent cations, with K+ being the best at stabilizing rG4 and Li+ having destabilizing effects, but alternate conformations of RNA are not greatly affected by these cations^37^. By comparing ThT fluorescence in the presence of rG4 stabilizing and destabilizing cations we strengthen the evidence that enhanced fluorescence is due to rG4 formation. TV2 induces significantly more fluorescence from ThT than TV2m, furthermore, the large difference in the fluorescence from the K+ and Li+ containing samples (Figure 6A) is indicative of rG4 formation being the underlying cause for the fluorescent enhancement by TV2. At lower RNA concentrations TV2 has the greatest difference in ThT fluorescence between K+ and Li+ samples but at higher RNA concentrations the difference in fluorescence tapers off, indicating the RNA may be forming rG4 at higher concentrations despite the unfavourable Li+ cation. TV2m has considerably less effect on ThT fluorescence, typically emitting 10-30% the amount of fluorescence compared to TV2 at a given concentration; together with a lack of significant difference between TV2m in K+ or Li+ containing samples this indicates that TV2m does not form a rG4.

**Figure 6.**
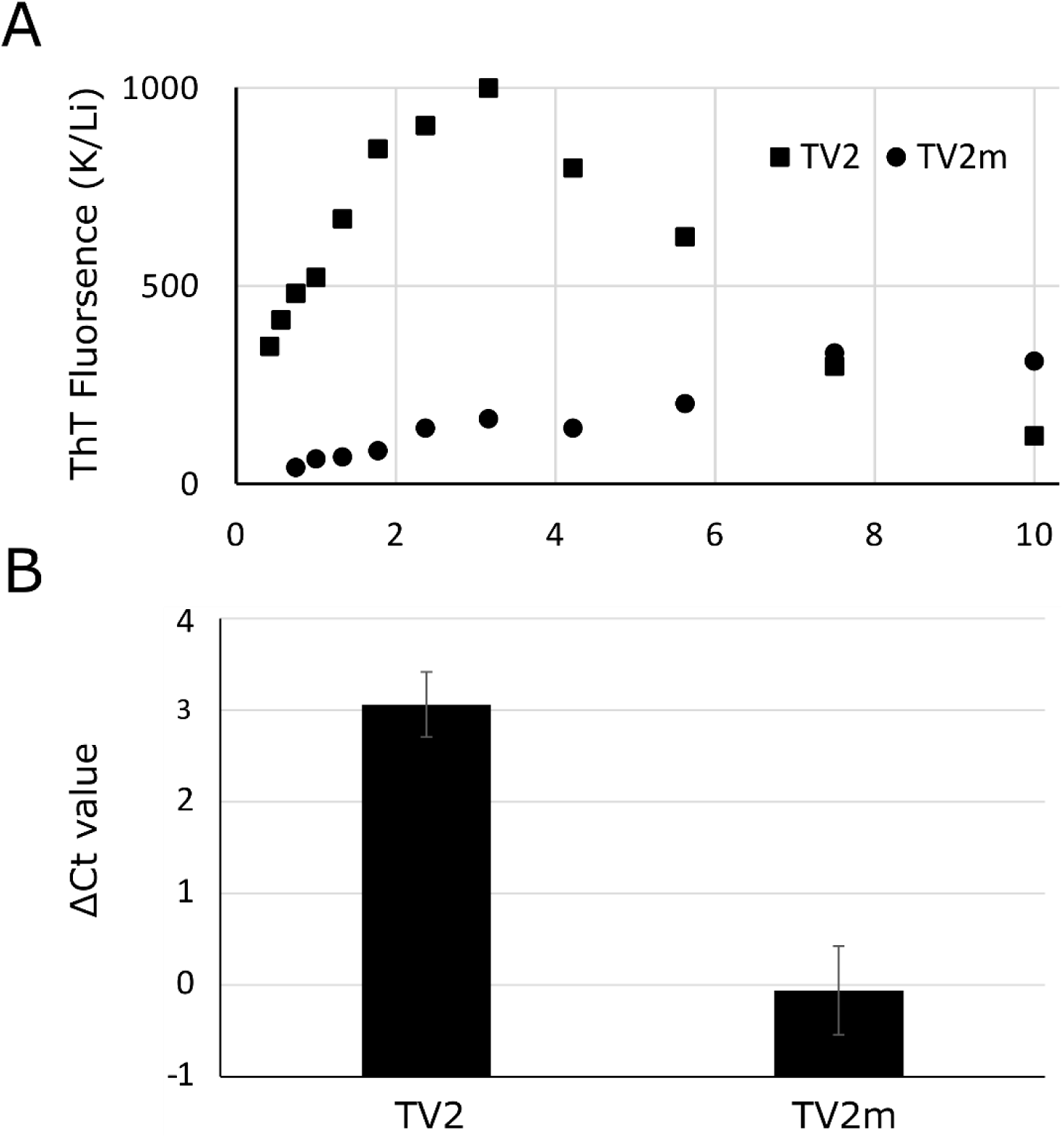
(A) Thioflavin T fluorescence enhancement by serial dilutions of TV2 and TV2m, the data is presented as the ratio of the normalized fluorescence ratio in K+ vs Li+ containing buffers. (B) Reverse transcriptase stop assay performed on TV2 and TV2m and quantified by qPCR, ΔCt values represent the difference between the Ct values for the full-length PCR product vs the 3’ terminal portion, which is the same between TV2 and TV2m and contains no GGG runs. Error bars represent the standard deviation between three reverse transcription reactions.

Finally, we used a modified reverse-transcriptase stop assay (rt-stop) to demonstrate structural differences between TV2 and TV2m. The principle of this assay is that a structured RNA will not be efficiently reverse transcribed and result in a mixture of full length and aborted transcripts. By comparing the amount of reverse transcriptase stopping TV2 and TV2m we can infer differences in their structure. Briefly, reverse transcription reactions were performed on equal amounts of both of the *in vitro* purified RNA at 37°C for 20 minutes. After purification of cDNA, qPCR was used to amplify the cDNA and the cycle at which the qPCR signal reached its critical threshold Ct was taken as an indicator of the efficiency of cDNA synthesis. We used two primer sets to probe the efficiency of reverse transcription of TV2 and TV2m RNA. One primer set spanned the entire RNA and the other primer set only amplified the 3’ region of the RNA after the last run of 3 consecutive guanines and where the two RNA share the same sequence. Reverse transcription products that stopped due to structures in the region of mRNA containing the G tracts would not be detected with the primer spanning the full UTR, whereas they could still be amplified by the second primer set making it a useful control for the amount of RNA input. For TV2m, both primer sets gave comparable Ct values, resulting in a ΔCt of 0.05 ± 0.48 cycles between them (Figure 6B). For the wild-type TV2 the Ct was reached 3.06 ± 0.35 cycles later with the primer set spanning the full UTR than the internal control primer set.

### Identification of rG4 location and evidence of rG4 unwinding by DDX21

To begin to characterize the rG4 forming sequence in the MAGED2 TV2 UTR we performed reverse transcription reactions on TV2 and TV2m using fluorescently labelled primers, this allowed us to visualize the rt-stop products on by denaturing PAGE (Figure 7A). The full length 234nt cDNA was observed in both TV2 and TV2m reverse transcription reactions, while a shorter, ∼115nt cDNA was only observed with TV2 (Figure 7B), This shorter cDNA is likely the result of a stalled reverse transcriptase that encountered significant secondary structure. From the length of this rt-stop product we can locate the position of the interfering structure in theTV2 RNA. The rt-stop is immediately prior to a sequence with high rG4 forming potential: (5’ggggctgggctgggttggg3’).

**Figure 7.**
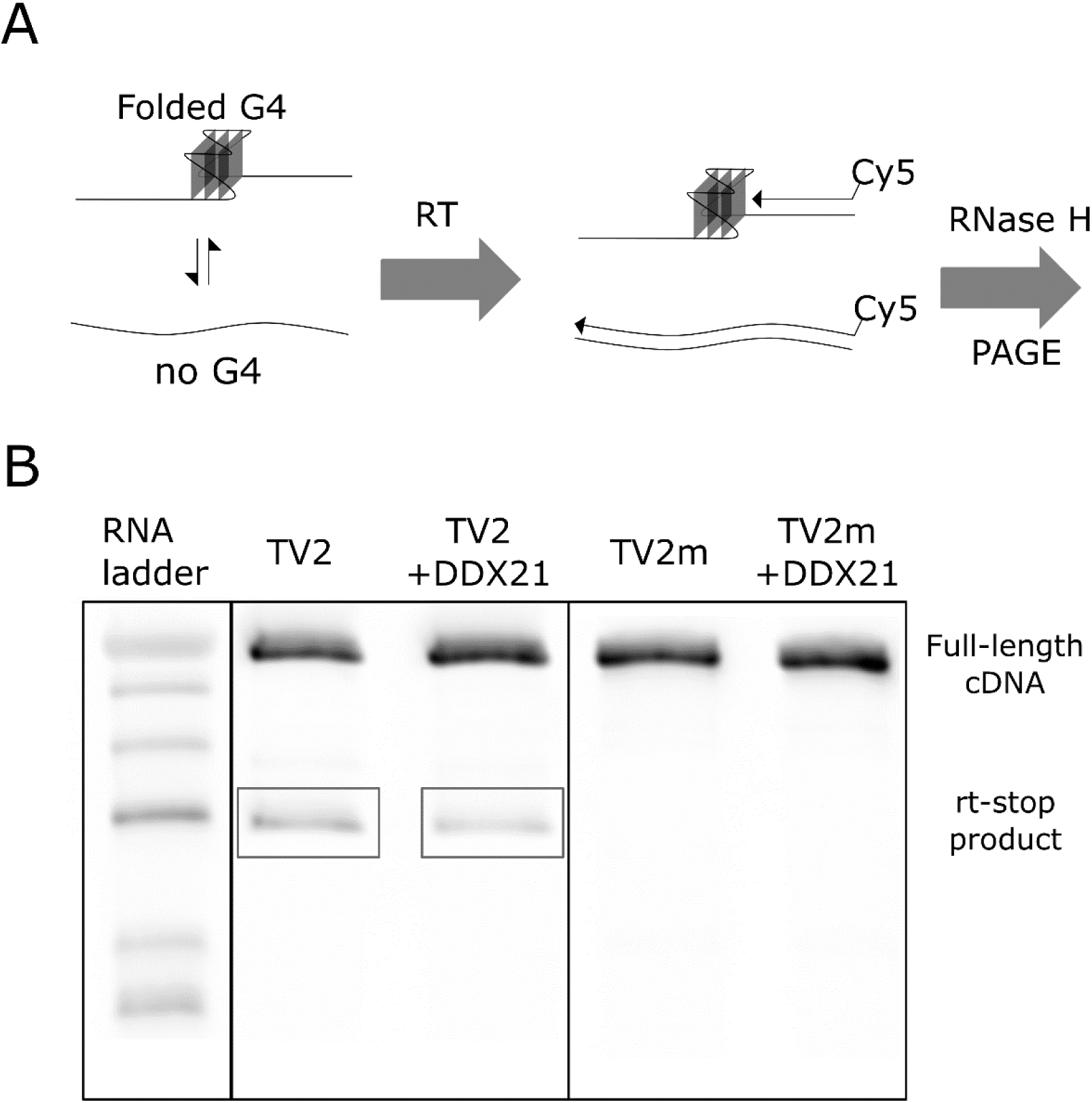
(A) Cartoon diagram showing the workflow of the rt-stop assay with fluorescent primers (B)Visualization of rt-stop products using fluorescently labelled primers for rt-reactions. The *in house* prepared RNA ladder (left) had 234nt, 200nt, 156nt, 119nt, 81nt and 69nt Cy5-labelled oligonucleotides. Full length cDNA products are observed for TV2 and TV2m adjacent to the 234nt ladder. Reverse transcription-stop products (boxed in grey) are observed only with the TV2 RNA template. Addition of DDX21 (56nM) to the rt reaction reduces the intensity of the rt-stop by 2-fold.

When recombinant purified DDX21 is added to the reverse transcription reaction with the TV2 RNA we observe a two-fold decrease in rt-stop products by densitometric analysis, remarkably consistent with the change in MAGED2 protein we see with DDX21 knock-down and M4 recovery (Figure 1A & Figure 3). This data indicates that DDX21 can unwind or destabilize the rG4 within TV2 enough to significantly alter the reverse-transcription efficiency.

## Discussion

Recently we discovered DDX21 to be an RNA rG4 binding protein that has the ability to destabilize rG4s^14^. Until now studies have focused on model RNA systems and there are no rG4s that DDX21 is known to interact with endogenously. A unique aspect of our study unique is the implementation of an RNA helicase protein that has been site-specifically mutated to have reduced affinity for RNA G-quadruplexes. By comparing differences in the whole-cell proteome in cell populations with either a wild type or mutant helicase protein we are able to discern differences that are due to the reduced affinity for RNA G-quadruplexes. We use this tool to identify, for the first time, potential biologically relevant G-quadruplex targets of the RNA helicase protein DDX21. This method could also be adapted to the study of other moonlighting proteins where the biochemical data exists to allow for specific mutations that only affect one of its activities.

From the list of proteins affected by replacement of wildtype DDX21 with M4 we validated MAGED2 by Western blot, a 2-fold decrease in the M4 DDX21 sample by MS is observed. MAGED2 has been shown to interact with p53 and affect transcription and protein levels of TRAIL-R2. It has been hypothesized that increased MAGED2 expression is a mechanism to selectively inactivate certain P53 targets (i.e. TRAIL-R2). Previous studies have shown that a decrease in MAGED2 protein results in upregulation of P53 and TRAIL-R2 at the mRNA and protein level. ^25,31^. We observed a similar effect in MCF-7 cells, revealing a pathway whereby WT DDX21 can facilitate MAGED2 translation which in turn blocks TRAIL-R2 expression. Interestingly, replacement of the WT DDX21 for the M4 mutant results in decreased MAGED2 protein and in turn an increase in TRAIL-R2 expression, indicating that G4 binding and unwinding is important for efficient translation of MAGED2 mRNA. This increase in TRAIL-R2 expression is significant enough to allow for TRAIL induced apoptosis in MCF-7 cells depleted of DDX21 or expressing only the M4 DDX21.

After determining that DDX21 preferentially binds the TV2 5’UTR, by DDX21-IP and rt-qPCR, we investigated a mutant version of TV2 in which all runs of GGG had been changed to GCG (TV2m). The TV2m 5’UTR was not impacted by DDX21 knock-down or recovery, whereas TV2 had decrease luciferase activity with depleted DDX21, M4 DDX21 and in the presence of rG4 stabilizing dye cPDS. Furthermore, *in vitro* transcribed and purified TV2m shows reduced affinity for DDX21 by EMSA and does not demonstrate the K+ sensitive ThT fluorescence response that TV2 does. These data provide strong evidence for rG4 formation in the MAGED2 TV2 5’ UTR.

Using an adapted reverse transcription stalling assay, modelled off the recently developed high-throughput rG4-seq method^24^, we demonstrate that TV2 contains a structural element, not present in TV2m, that impedes reverse transcription efficiency. By adding DDX21 into the reverse transcription reaction we turn this easy to perform rG4 detection method into a functional helicase assay. A major downfall of the current methods used to probe G4 helicase activity is the lack of sequence context surrounding the G4 structure. In this assay we can convincingly show rG4 disruption in the context of a full length 5’ UTR, in this case, where the rG4 is in the center of a 234 nucleotide RNA. In our opinion this is a significant advancement from the trap-based assays^38^, tetramolecular assays^39^ and nuclease sensitivity assays^14^ used previously. Together these data provide strong evidence that DDX21 can regulate MAGED2 translation via a rG4 in its 5’UTR.

Targeting of rG4 regulated pathways is an emerging area of therapeutic potential for fighting cancer^40,41^ and sensitization of TRAIL-resistant breast carcinoma to TRAIL mediated apoptosis is a current therapeutic target^27,29,30^. Our data underline the potential for the integration of rG4 therapeutics with an existing cancer treatment interest. Current research suggests that DDX21 has a unique specificity for certain rG4s that may be dependent on loop sequence^14,42^. Further work aims to identify the structure of the TV2 rG4 that DDX21 is recognizing, deciphering the specificity of DDX21 may potentially allow for the design of small molecules that specifically block its action.

## Materials and Methods

### Cell Culture and Reagents

The HEK293T cell line was a gift from Dr. Thomas Klonisch and the MCF-7 cell line was a gift from Dr. Spencer Gibson. Both cell lines were grown in Dulbecco’s Modified Eagle’s Medium from Thermo Fisher Scientific (Ottawa, Canada) supplemented with 10% Fetal Bovine Serum (Thermo Fisher Scientific). Gibco Trypsin/EDTA solution (Thermo Fisher Scientific) was used to detach the MCF-7 cells. Polyethyleneimine (Thermo Fisher Scientific) was used as the transfection reagent for vector DNA transfections

### DDX21 siRNA knock-down and recovery

Polyethyleneimine (Thermo Fisher Scientific) was used as the transfection reagent for vector DNA transfections. HEK293T cells were grown to 80% confluence in 150mm cell culture plates and transfected with empty pcDNA 3.1+ (negative control and knock-down samples) or siRNA resistant wt-DDX21 or m4-DDX21 expressing pcDNA3.1+. The following day the cells were split 1:9 and reverse transfected using transfectamine siRNA Max with either negative control siRNA (negative control sample) or DDX21 siRNA (Knock down and wild type and m4 recovery samples). 72 hours after reverse transfection the cells were collected in Phosphate Buffered Saline (Thermo Fisher Scientific) and rinsed twice. MCF-7 cells were treated the same way but cultured in 6-well dishes and only split 1:3 when reverse transfecting the siRNA. The remainder were lysed in ice cold RIPA buffer supplemented with 100x HALT proteinase cocktail (Thermo Fisher Scientific). Cells were vortexed and sonicated briefly before pelleting cell debris by centrifugation at 21,000 g. The protein concentration of the soluble portion was determined by Bradford assay and 50 µg of each sample was used for SDS-PAGE and Western blots.

### In-gel digestion

Protein lanes were visualized by Coomassie Blue staining prior to whole-lane excision. Each lane was subsequently cut into 15 equal bands, with each band corresponding to a region of the gel containing proteins of a distinct molecular weight range. Each of the gel fractions was subjected to in-gel tryptic digestion as previously described^41^; the resulting extracted peptides were then dried and suspended in 60 µL of 0.2% formic acid in 5% ACN.

### LC-MS/MS Analysis

Digested peptides were analyzed by LC-MS/MS using a ThermoScientific Easy nLC-1000 in tandem with a Q-Exactive Orbitrap mass spectrometer. Five microliters of each sample was subject to a 120 min gradient (0–45% buffer B; buffer A: 0.2% formic acid in 5% ACN; buffer B: 0.2% formic acid in ACN) on a 2 cm Acclaim 100 PepMap Nanoviper C18 trapping column in tandem with a New Objective PicoChip reverse-phase analytical LC column. For data-dependent analysis, the top 15 most abundant ions were analyzed for MS/MS analysis while +1 ions were excluded from MS/MS analysis. Additionally, a dynamic exclusion of 10 s was applied to prevent continued reanalysis of abundant peptides. For the analysis, a resolution of 35000 was used for full scans that ranged from 400 to 2000 m/z and a resolution of 17500 was used for MS/MS analysis.

For data analysis, raw data files corresponding to samples comprising an entire gel lane were grouped together and searched using Proteome Discoverer 1.4.1.14’s SEQUEST search algorithm using the reviewed, nonredundant *H. sapien* complete proteome retrieved from UniprotKB. Search parameters were identical to as previously reported^42^. During data processing, the “Precursor Ion Area Detector” node of Proteome Discoverer 1.4.1.14’s SEQUEST workflow editor was implemented to quantify the extracted ion chromatogram for each protein identified from the raw data. Searched results were filtered using a minimum of two medium confidence peptides per protein.

### Protein-RNA cop-immunoprecipitation and RT-qPCR

Native RNA immunoprecipitations were performed on one 150 mm dish of HEK293T cells as previously described^45^ using anti-DDX21 antibody (ProteinTech 15252-1-AP). RT-qPCR analysis was performed on Applied Biosystems StepOnePlus instrument with the RNA to Ct One-step RT-qPCR kit (Thermo Fisher Scientific) according to manufacturer’s instructions. To determine fold enrichment, RNA extracted from the total cell lysate using the GeneJET RNA purification kit (Thermo Fisher Scientific) was used to IP to serve as a reference sample. 25 ng of template RNA was used in all RT-qPCR reactions.

### Western blots

For each sample, 50 µg of protein in 1X SDS-PAGE load dye was loaded onto a 10% SDS-PAGE gel and ran at a continuous voltage (200V) for 45 minutes. Proteins were transferred to a 0.2 µm PVDF membrane (BioRad Laboratories) using the Trans-Blot Turbo transfer system (BioRad Laboratories). Membranes were blocked for 30 minutes in 5% milk powder dissolved in Tris Buffered Saline + Tween (TBST) (20 mM Tris pH 7.5, 150 mM NaCl, 0.1% Tween). The following antibodies were used: DDX21 (Novus NB100-1716), MAGED2 (ProteinTech 15252-1-AP), TRAIL-R2 (AbCam EPR19310, AB199357), CNOT3 (ProteinTech 11135-1-AP), XPO6 (ProteinTech 11408-1-AP), gapdh, tubulin, goat anti rabbit secondary, goat anti mouse secondary. DDX21, MAGED2 and TRAIL-R2 primary antibodies were incubated overnight at 4 degrees Celsius in TBST+ 5% milk, GAPDH and tubulin primary antibodies were incubated for 1 hour at room temperature in TBST+5% milk. Three sequential 5-minute washes with 5% TBS-T were followed by 1-hour incubation with secondary antibodies (1:10000) in TBST+5% milk and then 4 more 10-minute washes with TBS-T. Prior to imaging with the FluorChem Q system (Protein Simple), 1 ml of Illuminata Forte Western HRP substrate (EMD Millipore) was added to the blots. Due to weak signal from some antibodies a uniform adjustment to the brilliance was made using Inkscape V 0.91.

### Annexin V staining & TRAIL treatment

MCF-7 cells were treated as described in the DDX21 siRNA knock-down and recovery section. At 48 hours post knock-down TRAIL protein (SOURCE?) was added to the media to a final concentration of 50 ng/ml. Approximately 18 hours later the cells were washed with 1x trypsin and then incubated with 1x trypsin at 37 °C for 5 minutes. Detached cells were collected, washed with cold PBS and resuspended in 100 µl of cold 1X Annexin-binding buffer supplemented with AlexaFluor 488 annexin V (Thermo Fisher Scientific) and incubated in the dark for 15 minutes. Samples were then diluted to 500 µl with 1X Annexin-binding buffer and kept on ice. Annexin V binding was measured using the BD FACSCalibur platform and analyzed using the BD CellQuest Pro software (BD Biosciences, San Jose, CA), 10,000 cells from each sample were analyzed and each sample performed in biological triplicate.

### Luciferase assays

For luciferase assays, HEK293T cells were treated as described in the DDX21 siRNA knock-down and recovery section scaled down to a 6 well dish sample size. For carboxypyridostatin treated cells, cPDS was added to a concentration of 1uM in the DMEM at 48 hours post knock-down. Then the cells were transfected with 1ug of psiCheck-2 vector in 250 µl serum free DMEM with 1.5 µg of polyethyleneimine. After 24 hours the cells were collected in cold PBS. Half of the cells were used for RNA purification using the GeneJET RNA purification kit (Thermo Fisher Scientific) and the other half used in the luciferase assay. The Dual-Glo Luciferase Assay system (Promega, Madison Wi, USA) was used according to manufacturer instructions, luminescence was measured for 10 seconds using the (Pelka’s instrument) and each sample performed in at least biological triplicate. To control for transfection efficiency, luminescence from the hRluc protein, under control of the 5’UTR cloned into the NheI site was normalized to the signal from the hluc protein for each sample. Each sample was then normalized to the average value of the negative control samples.

### Cloning

The pcDNA3 vector was used to create a CMV promotor lacking nucleotides between 208 and 746, this was then cloned into the BglII and NheI sites of the psiCHECK-2 vector, removing the SV40 promotor. 5’UTRs of MAGED2 TV2 and TV3 were amplified by RT-PCR from RNA isolated from human testes tissue purchased from Takara Bio (Mountain View, CA). The 5’UTR of TV1 was amplified from a DNA fragment purchased from Genscript. Each TV was amplified using primers that contain restriction enzyme sites for NheI and cloned into the psiCHECK-2 vector. The cDNA that encodes the human DDX21 protein, isoform 1, was amplified by PCR using DNA primers that were designed to encode an NheI and NotI restriction enzyme sites and cloned into the pET28b expression vector.

### In vitro transcription and protein purification

The 5’UTRs of each transcript variant were transcribed and purified *in vitro* as previously described^46^ from linearized recombinant psiCHECK2 vector (Promega) that contained one of the three UTR sequences downstream of a T7 promoter. The vector was linearized with NcoI restriction enzyme and purified by phenol-chloroform extraction. *In vitro* transcription was carried out at 37°C for 3 hours. The transcription reaction was then stopped by addition of EDTA and T7 polymerase was removed by phenol-chloroform extraction. Residual phenol was removed using a DG10 desalting column (BioRad), from which the RNA was eluted in 20 mM Tris pH 7.5, 1 mM EDTA and either 100 mM KCl (TEK) or 100 mM LiCl (TELi). The RNA was then separated from the plasmid and remaining NTPs by FPLC purification using a Superdex 200 filtration column (GE Healthcare Life Sciences) using TEK or TELi buffer. FPLC fractions were then concentrated and stored at 4°C. RNAs were heated at 95°C and cooled on ice prior to use.

DDX21 was purified from LOBSTR *E. coli* BL21 (DE3) (Kerafast Inc, Boston, MA) cells transformed with DDX21 expressing pET28b. The stop codon from DDX21 cDNA was retained so that the recombinant protein only had the N-terminal 6His tag and thrombin cleavage site and not the C-terminal 6His tag. Cells were grown at 37°C in a shaker incubator to an optical density of 0.4 at A_600_, then induced with 0.3 mM IPTG and grown overnight at 18°C. Cells were pelleted, then resuspended in 20 mM Tris 7.5, 1 M NaCl, 10% glycerol, 1 mM DTT and 1 mM PMSF, and lysed by sonication. After centrifugation of lysate, recombinant DDX21 was purified from the soluble fraction using HisPur Cobalt Resin (Thermo-Fisher) and eluted using 200 mM imidazole. Elution fractions were dialyzed against 20 mM Tris pH 7.5, 300 mM NaCl, 10% glycerol, 2 mM DTT, and stored at 4°C.

### Electrophoretic Mobility Shift Assays

DDX21, or an equal volume of protein storage buffer (20 mM Tris pH 7.5, 300 mM NaCl, 10% glycerol, 2 mM DTT) was added to the binding reactions in a 1:1 serial dilution, with the first condition containing no DDX21. Binding reactions were performed in 50 mM Tris-acetate, pH 7.8, 100 mM KCl, 10 mM NaCl, 3 mM MgCl2, 70 mM glycine, 10% glycerol, 0.05 mg/mL bovine serum albumin (BSA) for 10 min at room temperature and resolved by native Tris-borate EDTA (TBE) 8% polyacrylamide gel electrophoresis (TBE-PAGE) for 150 minutes at 75V. The RNAs were then stained with the fluorescent nucleic acid dye SYBR Gold and imaged on a Fluorchem Q imaging system using the Cy2 excitation LEDs and emission filters (Protein Simple, San Jose CA).

### Thioflavin T assays

In a 96-well plate, the *in vitro* transcribed RNA was serially diluted 3:1 with 20 mM Tris pH 7.5, 1 mM EDTA and either 100mM KCl or 100 mM LiCl, from 10 µM to 0.4 µM. Thioflavin T was added with a final concentration of 1.5 µM. Samples incubated for 5 minutes at room temperature and fluorescence was measured with Applied Biosystems Step-One-Plus qPCR machine. Fluorescence data was extracted from the BLUE channel from the raw data output of the Step-One-Plus.

### Reverse Transcriptase Stop Assays

First strand synthesis of cDNA from 50ng of *in vitro* transcribed 5’UTR of TV2 and TV2m was performed in a 20µL reaction using 1µM reverse primer, 500µM dNTPs, 4µL 5X RT Buffer (250 mM Tris-HCl (pH 8.3 at 25 °C), 375 mM KCl, 15 mM MgCl2, 50 mM DTT) (Thermo Fisher), and 200U Maxima H-Minus RT (Thermo-Fisher). Reactions were carried out at 37°C for 60 minutes before heating at 85°C for 5 minutes to inactivate the RT enzyme. The cDNA was diluted 1:20 to the qPCR reaction using PowerUp Sybr Green Master Mix (Applied Biosystems), forward primers were added to a final concentration of 1µM. Forty cycles of qPCR were performed with 60°C annealing temperature and 30 second extension time using a Step-One-Plus qPCR machine (Applied Biosystems). CT values were determined using StepOne software. For visualization of rt stop products, first strand synthesis of cDNA from 1µg of *in vitro* transcribed TV2 and TV2m was performed in a 20µL reaction using 1µM Cy5 labeled primer, 500µM dNTPs, 4µL 5X RT Buffer (250 mM Tris-HCl (pH 8.3 at 25 °C), 375 mM KCl, 15 mM MgCl2, 50 mM DTT) (Thermo Fisher), and 200U Maxima H-Minus RT (Thermo-Fisher). Reactions were performed at 37°C for 60 minutes before heating at 85°C for 5 minutes to inactivate the RT. To remove DNA:RNA heteroduplex from the reactions, the cDNA samples were treated with RNAse H for 20 minutes at 37°C. After RNAse H digestion, cDNA was mixed with an equal volume of denaturing RNA load dye (95% formamide, 0.01% SDS, 0.5 mM EDTA, 0.025% (w/v) Orange G), heated at 95°C for 5 minutes, and resolved by TBE-urea PAGE with 12% acrylamide gels. Cy5 labelled RNA ladder was prepared as described previously^47^ with the addition of Cy5-labbeled TV2 for the additional 234nt band. Reverse transcription products and ladder were visualized with Cy5 excitation and emission filters.

