## Supplemental Information for "An RNA guanine quadruplex regulated pathway to TRAIL-sensitization by DDX21"

### Supplementary Information

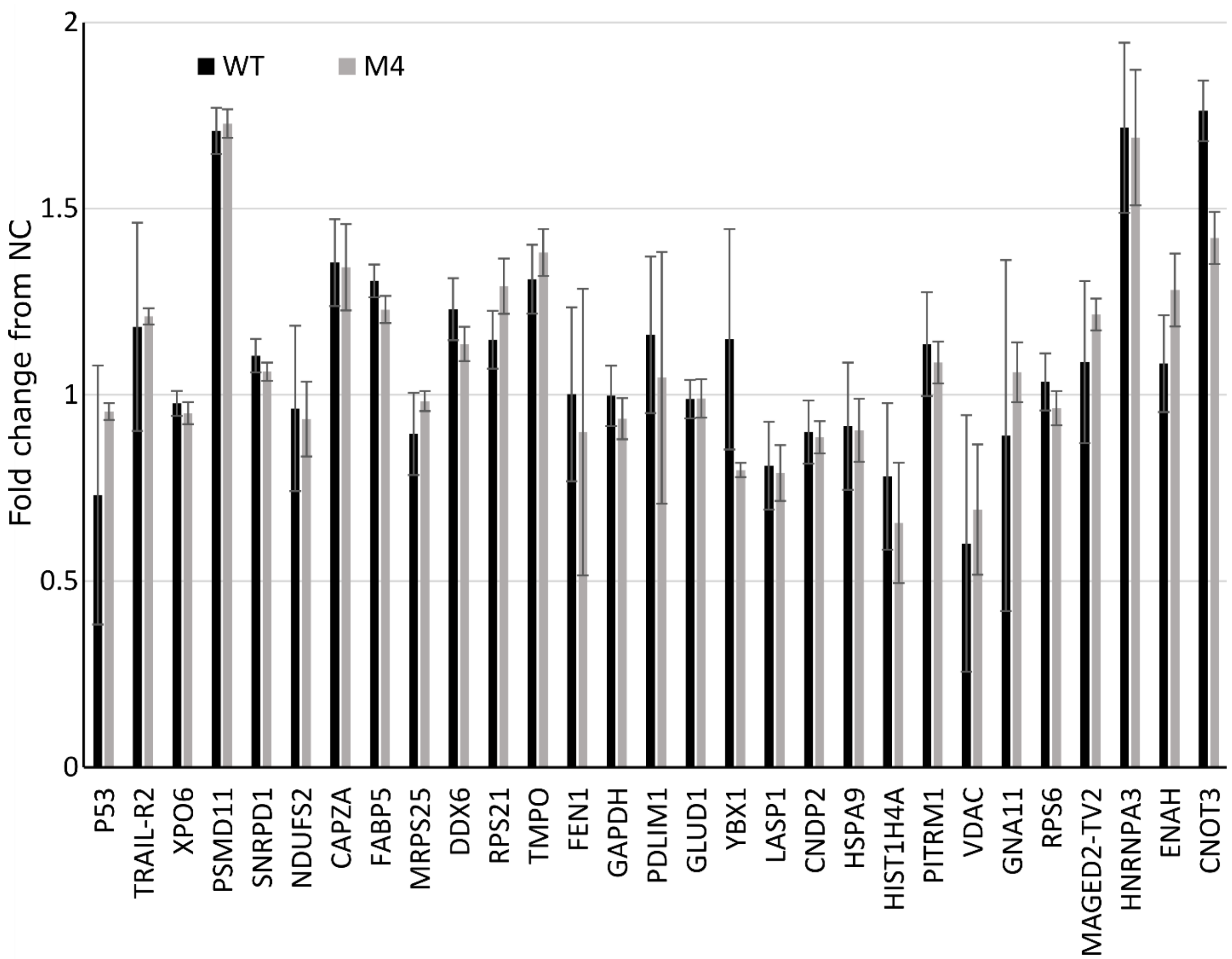


***Figure S1.*** RT-qPCR analysis comparing mRNA levels of target proteins between WT and M4 DDX21 recovery samples. Error bars represent the standard deviation between 3 biological replicates.

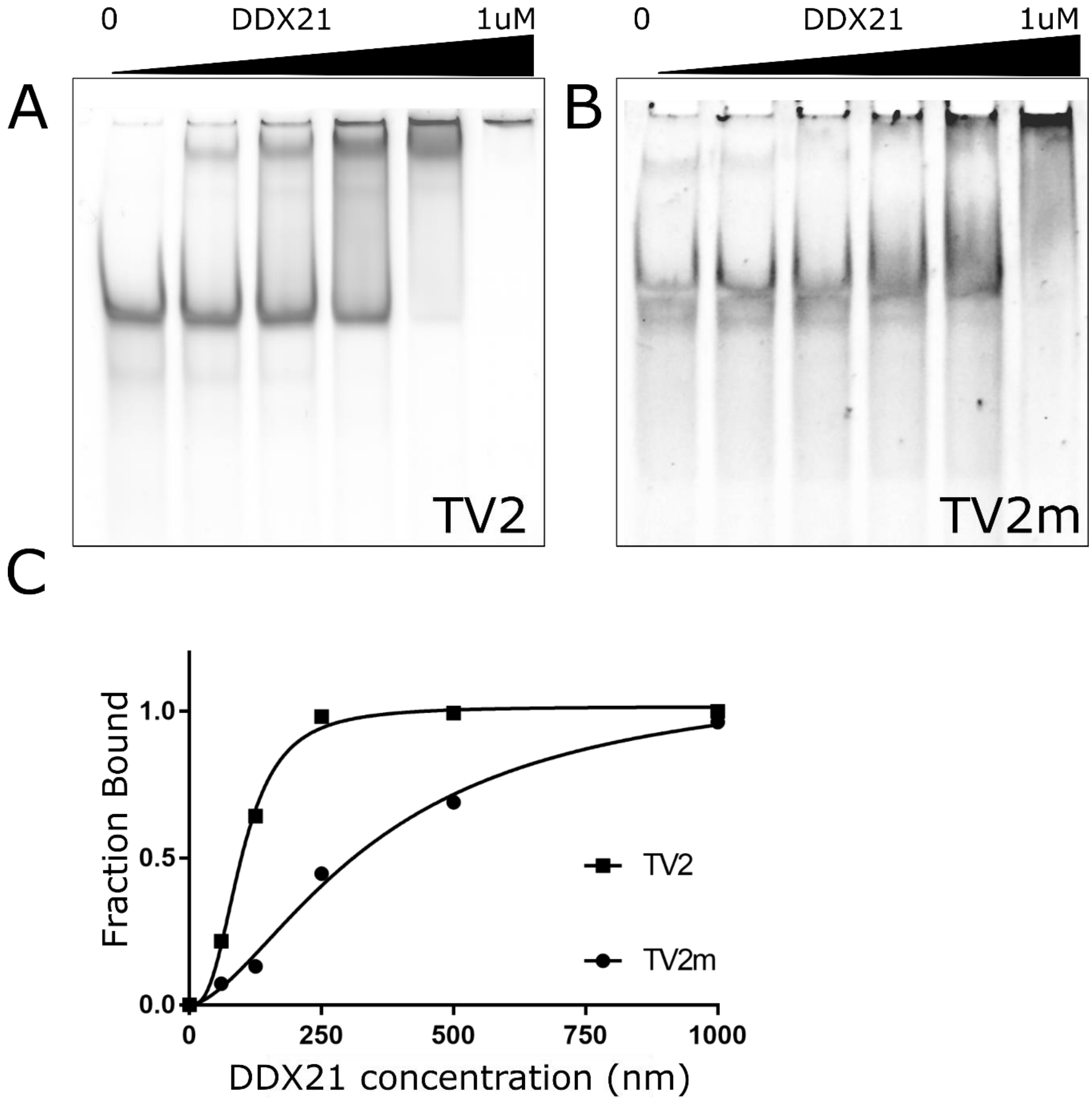


***Figure S2.*** EMSA of MAGED2 5’ UTRs, (A) TV2, (B) TV2m with a serial (1:1) dilution of DDX21. Fitting curves (C) determined in Prism from a Specific binding model with Hill slope from densitometric analysis of EMSA images.
